# Development of nuclear microsatellite markers in Yerba mate (*Ilex paraguariensis* St. Hil., Aquifoliaceae Bartl.) from whole-genome sequence data

**DOI:** 10.1101/2022.12.21.521502

**Authors:** Carolina Tassano, Rodrigo A. Olano, Paola Gaiero, Magdalena Vaio, Pablo R. Speranza

## Abstract

*Ilex paragariensis* St. Hil. (*yerba mate*) (Aquifoliaceae Bartl.) is a plant species with great economic and cultural importance because its leaves are processed and ground to make infusions like mate or tereré. The species is distributed in a continuous area that includes southern Brazil, and part of Paraguay and Argentina. Uruguay represents the southern distribution limit of the species, where small populations can be found as part of riparian forests. Although there are previous reports of molecular markers for this and other species in the genus, the available markers were not informative enough to represent the intra and interpopulation genetic diversity in these marginal populations. In this study, we developed highly informative nuclear polymorphic microsatellite markers to optimize genetic studies in *I. paraguariensis*. Markers were identified in contigs from the genome sequence of two individuals and then tested for amplification and polymorphism in a diverse panel and at population level. Markers which passed these tests detected levels of heterozygosity similar to those reported for Brazilian populations and great diversity within populations from Uruguay. This set of markers were successfully multiplexed, substantially reducing the costs of the analysis. In combination with previously reported nuclear and plastid markers, they can be used to evaluate the genetic diversity of rear edge populations, identify genotypes for paternity studies and provide relevant information for the conservation and management of germplasm.

## INTRODUCTION

*Ilex paraguariensis* St. Hil. (*yerba mate*) is a plant species with great economic and socio-cultural importance. The drinks called mate and tereré, consumed mostly in Uruguay, Brazil, Argentina and Paraguay, are made with its leaves. *I. paraguariensis* is a perennial subtropical tree, distributed in southern Brazil, part of Paraguay and Argentina. Uruguay represents its southern distribution limit, where small populations are found in riparian forests (Grela 2004; Hernández 2019).

Molecular markers previously reported for the genus *Ilex* were useful but had some limitations for the study of genetic diversity in marginal populations (Gottlieb et al. 2011; Cascales et al. 2014; Talavera 2021). Markers developed for *I. leucoclada* (Torimaru et al. 2004) and *I. paraguariensis* (Pereira et al. 2013) showed reduced genetic variation in populations from Uruguay, which otherwise are expected to show high differentiation and alleles not found in the central area of the distribution (Hampe and Petit 2005). Moreover, plastid microsatellite markers were mostly monomorphic or showed very low polymorphism (Cascales et al. 2014; Hernández 2019; Talavera 2021). Therefore, SSR markers specifically designed to maximize the representation of diversity in marginal populations, in combination with those previously reported, will be useful to evaluate genetic diversity of rear edge populations, for paternity studies and germplasm conservation and management. In this study, we developed highly informative nuclear polymorphic microsatellite markers to optimize genetic studies in *I. paraguariensis*.

## EXPERIMENTAL

### Plant material

To characterize the markers, we used a diverse panel from 12 populations of *I. paraguariensis* from Uruguay and one from Paraguay (one individual per population, Table 1). In order to characterize a subset of these markers at the population level, a sample of 15 individuals from three populations was used (Table 2). Leaves were collected and dried in silica. DNA extraction was performed using a standard protocol (Doyle and Doyle 1987).

**Table 1.**
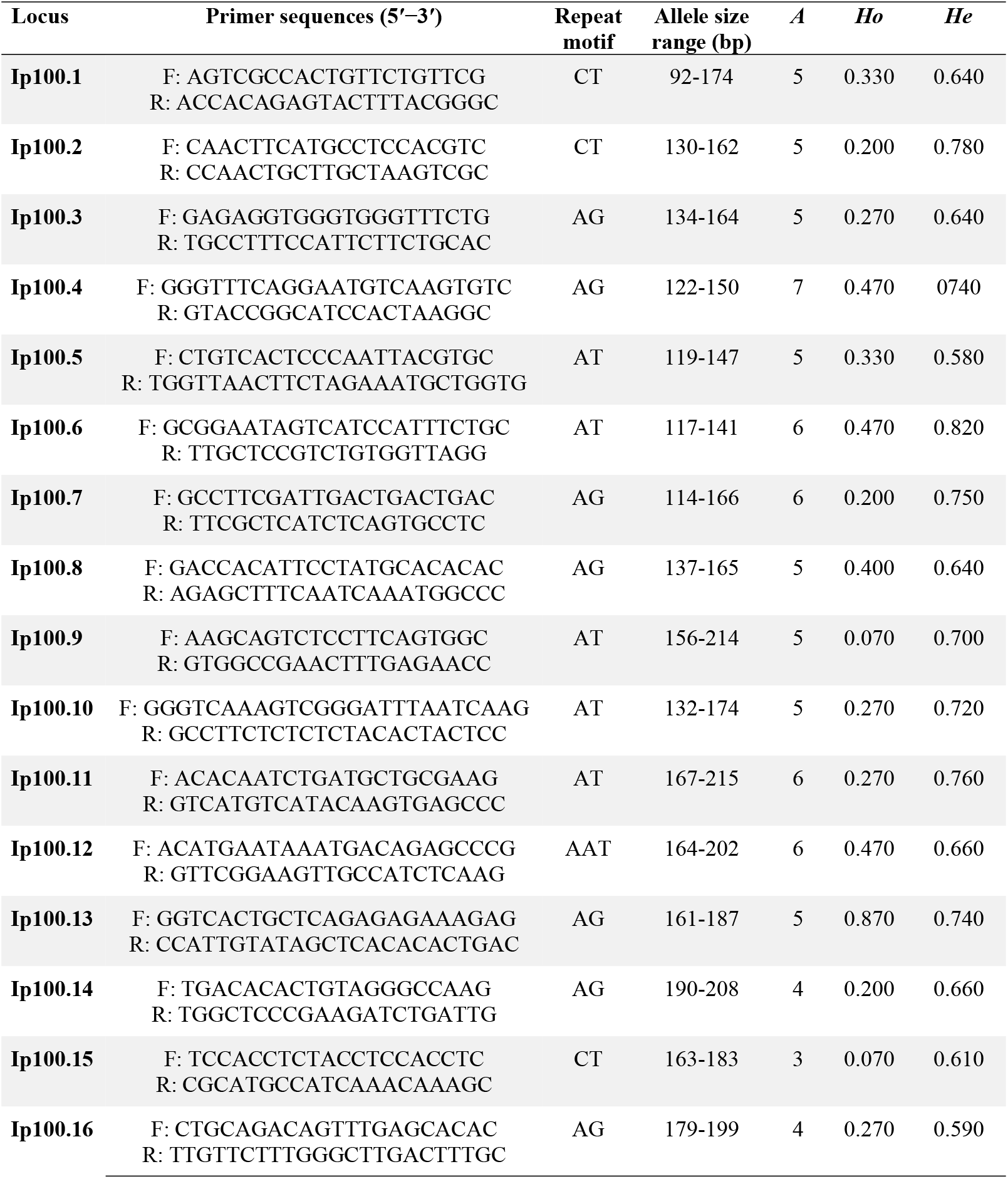
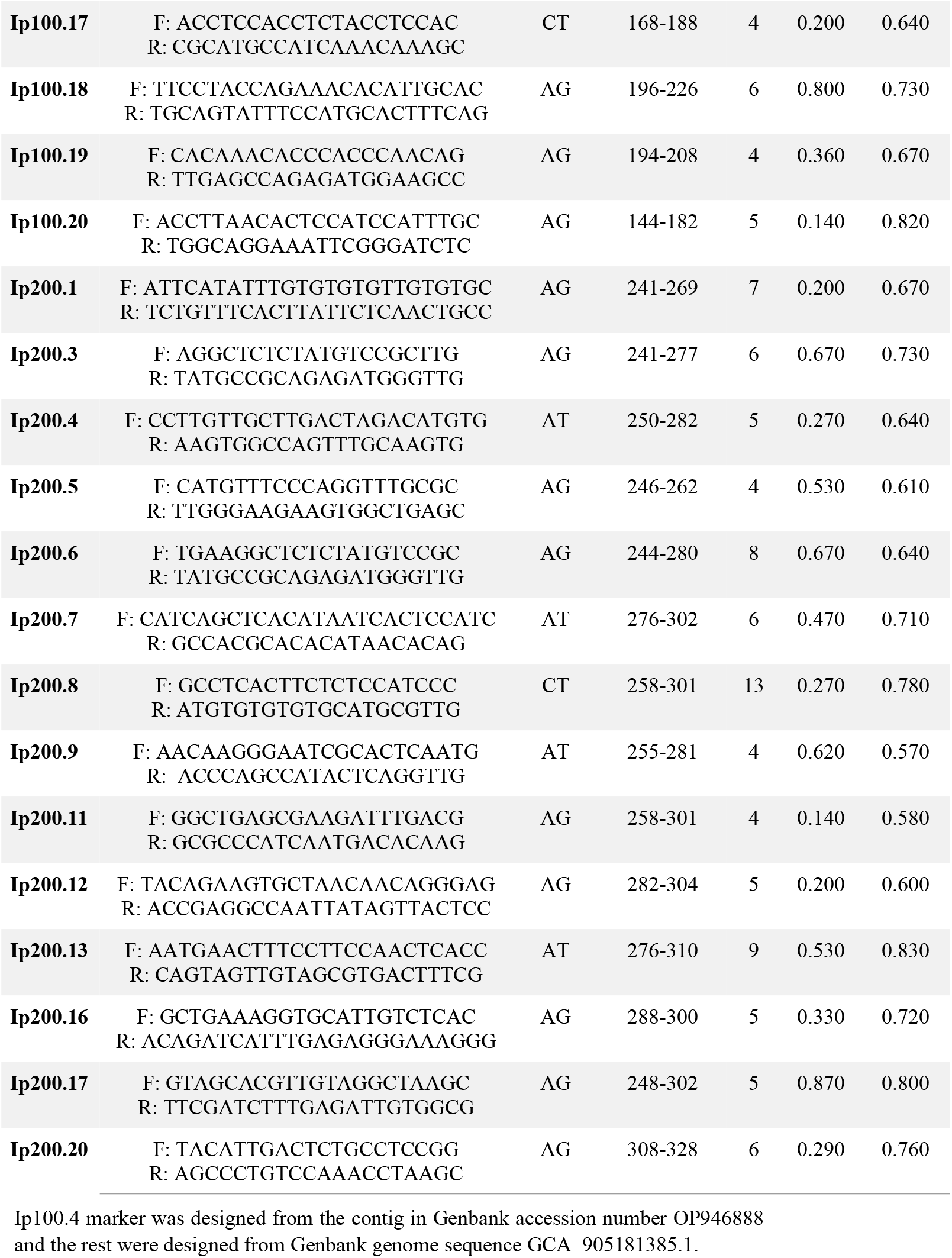
Characteristics of microsatellite markers developed for *Ilex paraguariensis* from genomic data, including forward (F) and reverse (R) primer sequences, repeat motif, observed allele size range; *A* = number of alleles; *Ho* = observed heterozygosity, *He* = expected heterozygosity. Annealing temperature for all primer pairs was 60 °C

**TABLE 2.** Genetic diversity properties of 7 polymorphic microsatellite markers developed for *Ilex paraguariensis* characterized in three populations from Uruguay, n, sample size for each population. Parameters detailed for each marker are *A* = number of alleles; *Ho* = observed heterozygosity, *He* = expected heterozygosity, *HWE:* χ^2^ values for the test of Hardy–Weinberg equilibrium, *Fst* = genetic differentiation coefficient.

| Loci | Population 1<br>(n=14) |  |  |  |  |  | Population 2<br>(n=15) |  |  |  |  |  | Population 3<br>(n=15) |  |  |  |  |  |
| --- | --- | --- | --- | --- | --- | --- | --- | --- | --- | --- | --- | --- | --- | --- | --- | --- | --- | --- |
|  | *A* | *Ho* | *He* | *HWE* |  | *Fst* | *A* | *Ho* | *He* | *HWE* |  | *Fst* | *A* | *Ho* | *He* | *HWE* |  | *Fst* |
| **Ip100.2** | 3 | 0.429 | 0.406 | 1.238 | ns | -0,057 | 4 | 0.400 | 0.391 | 12.267 | ns | -0,023 | 5 | 0.667 | 0.593 | 16.038 | ns | -0,124 |
| **Ip100.6** | 4 | 0.538 | 0.615 | 8.343 | ns | 0,067 | 3 | 0.500 | 0.622 | 7.202 | ns | 0,196 | 3 | 0.333 | 0.531 | 16.445 | *** | 0,372 |
| **Ip100.11** | 3 | 0.429 | 0.523 | 4.300 | ns | 0,180 | 1 | 0.000 | 0.000 | 14.000 | *** | 0.000 | 3 | 0.600 | 0.464 | 4.945 | ns | -0,292 |
| **Ip100.20** | 7 | 0.462 | 0.577 | 35.902 | * | 0,200 | 3 | 0.000 | 0.611 | 12.000 | ** | 1.000 | 2 | 0.400 | 0.444 | 0.150 | ns | 0,100 |
| **Ip200.3** | 5 | 0.643 | 0.740 | 16.918 | ** | 0,131 | 6 | 0.933 | 0.713 | 27.975 | * | -0,308 | 5 | 0.933 | 0.749 | 14.225 | ns | -0,246 |
| **Ip200.8** | 2 | 0.214 | 0.191 | 0.202 | ns | -0,120 | 5 | 0.533 | 0.660 | 32.417 | *** | 0,192 | 4 | 0.800 | 0.640 | 4.289 | ns | -0,250 |
| **Ip200.17** | 4 | 0.500 | 0.559 | 11.375 | ns | 0,105 | 7 | 0.667 | 0.631 | 7.252 | ns | -0,056 | 7 | 0.600 | 0.558 | 2.792 | ns | -0,076 |

Statistical deviation from HWE is indicated as: ns=not significant, \* P<0.05, \*\* P<0.01, \*\*\* P<0.001

### Sequences

Intact genomic DNA from one individual plant was used for library prep and low pass whole-genome sequencing using a DNBseq Illumina platform (150 bp reads, pair-end sequencing) at BGI Genomics (Tai Po, Hong Kong). A total of 1.41 Gbp of sequences were assembled into contigs using SOAPdenovo2 (see Suppl. file 1 for config file) (Luo et al., 2012). Only marker Ip100.4 was selected from one of these contigs (Table 1). The rest of the markers were developed from whole-genome sequences for *I. paraguariensis* assembled and deposited in GenBank by Sosa & Modenutti (2021). In both cases, microsatellite-like nuclear sequences were identified with Phobos 3.3.11 and primers complementary to their flanking regions were designed using Primer3 (Rozen & Skaletsky, 2000), both in Geneious 9.0 (Kearse et al., 2012). Sequences containing perfect repeats at least 15-units long were selected for primer design.

### Primer design

Primers were designed to obtain two sets of product sizes, 100-200 bp and 250-300 bp, all with a standardized annealing temperature of 60 °C. We selected primers with no repetitive sequences and no neighbouring microsatellites within the flanking region. A total of 40 primer pairs were ordered from the Custom DNA oligo synthesis service at Macrogen, South Korea (https://dna.macrogen.com/). Forward primers were extended with one of the following sequences to match a set of oligonucleotides labelled with FAM, VIC, NED and PET, respectively: 5’-AATACAACGCGATCGACTCC-3’; 5’-AATCCCCACACAAACACACC-3’; 5’-TCCCCTTTCAAACCTAATGG-3’; 5’-TGATCTTGAGAAGGCATCCA-3’.

### Amplification

Amplifications were performed in a Vertity thermal cycler (Applied Biosystems) and amplification products were run in a ABI3500 XL sequencer (Applied Biosystems). PCR cycling conditions consisted of an initial denaturation at 95°C for 15 min; 35 cycles of denaturation at 95°C for 30 s, annealing between 53°C to 60°C for 90 s, and extension at 72°C for 30 s; and a final extension cycle at 60°C for 30 min. For the population analysis, seven of the most informative markers were combined in a multiplex reaction. PCR multiplex amplifications contained 10 ng of genomic DNA, 0.75 uM of each forward primer and 3 uM of each reverse primer (for product sizes 100-200 bp) or 1 uM of each forward primer and 4 uM of each reverse primer (for product sizes of 250-300 bp), 1,25 μl 2X of Platinum Multiplex PCR Master Mix and 0,3 mL GC Enhancer (Applied Biosistems), 10X primer mix and ultrapure water. All analyses were performed at Genexa (https://www.genexa.com.uy/).

### Data analysis

Electropherograms were analyzed individually with PeakScanner 1.0 © (Applied Biosystems, 2006). Data were analyzed with GenAlEx 6.5 of Microsoft Excel (Peakall and Smouse, 2006).

## DISCUSSION

Most markers contain AG (55.8%), AT (26.4%) and CT (1.7%) dinucleotide repeats, and only one marker is an AAT trinucleotide. All of the 34 successfully amplified markers were polymorphic (Table 1). The number of alleles in the diverse panel ranged between 3 and 13 (average 5.5). Allele sizes ranged from 92bp to 226bp and between 241 and 328 bp. Non-overlapping size ranges allowed easy scoring of at least two loci labelled with the same fluorescent dye.

All seven loci included in the population analysis were polymorphic within all three populations. The number of alleles per locus ranged from 2 to 7 (average 4.0 alleles per population (Table 2). The level of Ho and He ranged from 0 to 0.933 (average 0,504) and from 0,191 to 0,749 (average 0,534) respectively (Table 2). Significant deviations from HWE based on Fisher’s exact test (P < 0.05) were detected for two loci in populations 1, four loci in population 2 and one loci in population 3 (Table 2).

Similar or superior levels of heterozygosity were detected in species with biology similar to *I. paraguariensis* (Sosa et al. 2010, Mantiquilla et al. 2021). Our markers allowed detecting levels of heterozygosity similar to those reported for Brazilian populations by Pereira et al. (2013) and a diversity within the Uruguayan populations greater than that reported by Cascales et al. (2014). These markers showed a high fixation of genetic diversity between populations (Table 2). Additionally, because all primers had one Tm value, and two size ranges, these markers were successfully multiplexed and analysed, substantially reducing the costs of analysis. Our results confirm the reliability of these markers to evaluate genetic diversity, population structure and conservation status in *Ilex paraguariensis.*

## Acknowledgements

This research was funded by Comisión Sectorial de Investigación Científica from the University of the Republic (22320200200217UD and 22320200200248UD) and by Agencia Nacional de Investigación e Innovación (ANII, grant number FCE_1_2021_1_166709). In addition, CT and RAO thank ANII and the Comisión Académica de Posgrados from the University of the Republic (CAP-Udelar) for scholarships funding. We also thank Pablo Hernandez and the land owners for collaborating in the field collections and providing plant material.

