## Supplemental Table 1 for "Development of nuclear microsatellite markers in Yerba mate (*Ilex paraguariensis* St. Hil., Aquifoliaceae Bartl.) from whole-genome sequence data"

| Locus | Primer sequences (5′−3′)  Forward | Primer sequences (5′−3′)  Reverse |
| --- | --- | --- |
| Ip100.1 | AGTCGCCACTGTTCTGTTCG | ACCACAGAGTACTTTACGGGC |
| Ip100.2 | CAACTTCATGCCTCCACGTC | CCAACTGCTTGCTAAGTCGC |
| Ip100.3 | GAGAGGTGGGTGGGTTTCTG | TGCCTTTCCATTCTTCTGCAC |
| Ip100.4 | GGGTTTCAGGAATGTCAAGTGTC | GTACCGGCATCCACTAAGGC |
| Ip100.5 | CTGTCACTCCCAATTACGTGC | TGGTTAACTTCTAGAAATGCTGGTG |
| Ip100.6 | GCGGAATAGTCATCCATTTCTGC | TTGCTCCGTCTGTGGTTAGG |
| Ip100.7 | GCCTTCGATTGACTGACTGAC | TTCGCTCATCTCAGTGCCTC |
| Ip100.8 | GACCACATTCCTATGCACACAC | AGAGCTTTCAATCAAATGGCCC |
| Ip100.9 | AAGCAGTCTCCTTCAGTGGC | GTGGCCGAACTTTGAGAACC |
| Ip100.10 | GGGTCAAAGTCGGGATTTAATCAAG | GCCTTCTCTCTCTACACTACTCC |
| Ip100.11 | ACACAATCTGATGCTGCGAAG | GTCATGTCATACAAGTGAGCCC |
| Ip100.12 | ACATGAATAAATGACAGAGCCCG | GTTCGGAAGTTGCCATCTCAAG |
| Ip100.13 | GGTCACTGCTCAGAGAGAAAGAG | CCATTGTATAGCTCACACACTGAC |
| Ip100.14 | TGACACACTGTAGGGCCAAG | TGGCTCCCGAAGATCTGATTG |
| Ip100.15 | TCCACCTCTACCTCCACCTC | CGCATGCCATCAAACAAAGC |
| Ip100.16 | CTGCAGACAGTTTGAGCACAC | TTGTTCTTTGGGCTTGACTTTGC |
| Ip100.17 | ACCTCCACCTCTACCTCCAC | CGCATGCCATCAAACAAAGC |
| Ip100.18 | TTCCTACCAGAAACACATTGCAC | TGCAGTATTTCCATGCACTTTCAG |
| Ip100.19 | CACAAACACCCACCCAACAG | TTGAGCCAGAGATGGAAGCC |
| Ip100.20 | ACCTTAACACTCCATCCATTTGC | TGGCAGGAAATTCGGGATCTC |
| Ip200.1 | ATTCATATTTGTGTGTGTTGTGTGC | TCTGTTTCACTTATTCTCAACTGCC |
| Ip200.3 | AGGCTCTCTATGTCCGCTTG | TATGCCGCAGAGATGGGTTG |
| Ip200.4 | CCTTGTTGCTTGACTAGACATGTG | AAGTGGCCAGTTTGCAAGTG |
| Ip200.5 | CATGTTTCCCAGGTTTGCGC | TTGGGAAGAAGTGGCTGAGC |
| Ip200.6 | TGAAGGCTCTCTATGTCCGC | TATGCCGCAGAGATGGGTTG |
| Ip200.7 | CATCAGCTCACATAATCACTCCATC | GCCACGCACACATAACACAG |
| Ip200.8 | GCCTCACTTCTCTCCATCCC | ATGTGTGTGTGCATGCGTTG |
| Ip200.9 | AACAAGGGAATCGCACTCAATG | ACCCAGCCATACTCAGGTTG |
| Ip200.11 | GGCTGAGCGAAGATTTGACG | GCGCCCATCAATGACACAAG |
| Ip200.12 | TACAGAAGTGCTAACAACAGGGAG | ACCGAGGCCAATTATAGTTACTCC |
| Ip200.13 | AATGAACTTTCCTTCCAACTCACC | CAGTAGTTGTAGCGTGACTTTCG |
| Ip200.16 | GCTGAAAGGTGCATTGTCTCAC | ACAGATCATTTGAGAGGGAAAGGG |
| Ip200.17 | GTAGCACGTTGTAGGCTAAGC | TTCGATCTTTGAGATTGTGGCG |
| Ip200.20 | TACATTGACTCTGCCTCCGG | AGCCCTGTCCAAACCTAAGC |

**Supplementary table**1. Forward (F) and reverse (R) primers sequences for each marker locus designed for *Ilex paraguariensis* from whole-genome sequence data.
