## Supplemental file 2 for "Development of nuclear microsatellite markers in Yerba mate (*Ilex paraguariensis* St. Hil., Aquifoliaceae Bartl.) from whole-genome sequence data"

#maximal read length
max_rd_len=150
[LIB]
#average insert size
avg_ins=500
#if sequence needs to be reversed
reverse_seq=0
#in which part(s) the reads are used
asm_flags=3
#use only first 100 bps of each read
rd_len_cutoff=150
#in which order the reads are used while scaffolding
rank=1
# cutoff of pair number for a reliable connection (at least 3 for short insert size)
pair_num_cutoff=3
#minimum aligned length to contigs for a reliable read location (at least 32 for short insert size)
map_len=32
#a pair of fastq file, read 1 file should always be followed by read 2 file
q1=/home/pgaiero/Ipar/assembly2/V300046929_L3_AE59379111-602_1.fq
q2=/home/pgaiero/Ipar/assembly2/V300046929_L3_AE59379111-602_2.fq
